# GeneSelectR: An R Package Workflow for Enhanced Feature Selection from RNA Sequencing Data

**DOI:** 10.1101/2024.01.22.576646

**Authors:** Damir Zhakparov, Kathleen Moriarty, Damian Roqueiro, Katja Baerenfaller

## Abstract

**Motivation:** High-dimensional Bulk RNA sequencing (RNAseq) datasets pose a considerable challenge in identifying biologically relevant features for downstream analyses and data mining efforts. The standard approach involves differential gene expression (DGE) analysis, but its effectiveness can be limited depending on the data due to its univariate nature. In complex datasets, an alternative approach involves employing a variety of machine learning (ML) tools, which attempt to understand non-linear relationships between features and focus on generalizability rather than statistical significance. This approach will result in the generation of multiple feature lists, which might exhibit similarities in terms of classification performance metrics. Therefore, there is an urgent need for a cohesive workflow that seamlessly integrates robust feature selection using diverse ML methods while also evaluating the biological relevance of the resulting feature lists. This combined approach would enable the prioritization of the best-performing list, considering both sets of criteria.

**Results:** We introduce GeneSelectR, an open-source R package that innovatively combines ML and bioinformatic data mining approaches for enhanced feature selection. With GeneSelectR, features can be selected from a normalized RNAseq dataset with a variety of ML methods and user-defined parameters. This is followed by an assessment of their biological relevance with Gene Ontology (GO) enrichment analysis, along with a semantic similarity analysis of the resulting GO terms. Additionally, similarity coefficients and fractions of the GO terms of interest are calculated. With this, GeneSelectR optimizes ML performance and rigorously assesses the biological relevance of the various lists, offering a means to prioritize feature lists with regard to the biological question. When applied to the TCGA-BRCA dataset, the GeneSelectR workflow generated several feature lists using different ML methods and a DGE analysis. By leveraging the various functions in GeneSelectR, the different lists could be evaluated based on both ML performance and biological relevance. This comprehensive evaluation facilitated the selection of the best-performing list, which exhibited both strong machine learning performance and high relevance to the biological question while maintaining a manageable number of highly specific features.

**Availability:** The package is available on CRAN. To install it, run: install.packages(‘GeneSelectR’)

**Supplementary information:** Supplementary data are available at *Bioinformatics* online.

## 1 Introduction

With bulk RNA sequencing (RNAseq) thousands of transcripts can be profiled in a high-throughput manner. The resulting datasets are typically high-dimensional and show an unfortunate imbalance between the number of features and samples, posing a challenge for the subsequent identification of significant features. The standard approach to select differentially expressed transcripts is statistical differential gene expression (DGE) analysis. Despite its effectiveness, DGE analysis has limitations, particularly its univariate nature, which becomes apparent in scenarios involving an experimental design with multiple variables, where the test variable is not the primary driver of variance in the dataset. In such cases, the statistical analysis may often yield a limited number of significant features after correcting for multiple testing, leading to reduced sensitivity in detecting genuine differences and an increased risk of type II errors. If the dataset is large enough, feature selection using machine learning (ML) is a viable option here to account for complex nonlinear relationships between genes. This procedure involves finding a subset of features associated with the target variable of the experimental design, capturing interactions between features that would otherwise remain hidden. When various Machine Learning (ML) methods are applied to an RNASeq dataset to identify significant features, they generate different lists of selected features. The evaluation of these lists primarily relies on ML classification performance metrics such as accuracy or precision (Chiesa, Colombo, and Piacentini 2018; Dag et al. 2022). However, particularly in complex datasets, due to the distinct working principles of various ML methods, they can select considerably different feature lists while still demonstrating similar classification performance metrics. Therefore, relying solely on these metrics does not guide the selection of the best list. Here, we introduce GeneSelectR, a novel workflow that combines the extensive ML capabilities of the sklearn Python package with the comprehensive functionality of bioinformatics R libraries with which the biological relevance of the feature lists can be assessed in addition.

## 2 Package Description

In this description, we outline the core functionalities of the GeneSelectR package. For more detailed information, please consult the vignette and Supplementary Material, which provides a step-by-step guide to analyzing the TCGA-BRCA dataset in R markdown. The input RNA sequencing data and the output files of the TCGA-BRCA use case are available at https://github.com/dzhakparov/GeneSelectR_benchmarking.

### 2.1 Input Data Requirements

GeneSelectR requires a vector containing sample labels and a data frame of dataset-wide normalized RNAseq read counts. The counts should be preprocessed through either DESeq2 or edgeR, ideally already mitigating for batch effects or other sources of variation and applying a variance-stabilizing transformation (Robinson, McCarthy and Smyth 2010; Love, Huber and Anders 2014).

### 2.2 Feature Selection Procedure

To make the GeneSelectR feature selection process user-friendly, we implemented the ML capabilities of the sklearn Python library via the reticulate R package, which combines it with the diverse R bioinformatics toolkit (Pedregosa 2012; Ushey K, Allaire J and Tang Y 2023). In addition, we containerized it with Docker so that it can be run in an isolated environment to avoid dependency conflicts and to minimize the need for frequent intermediate outputs and data transfers.

The GeneSelectR workflow, depicted in Figure 1, starts with data input, followed by preprocessing steps such as removing low variance features and scaling the data. Subsequently, the four default feature selection methods, namely Boruta, RandomForest, Univariate feature selection, and SelectFromModel with Logistic Regression (L1 penalty), are applied. Each of these methods undergoes the following steps:

**Fig 1.**
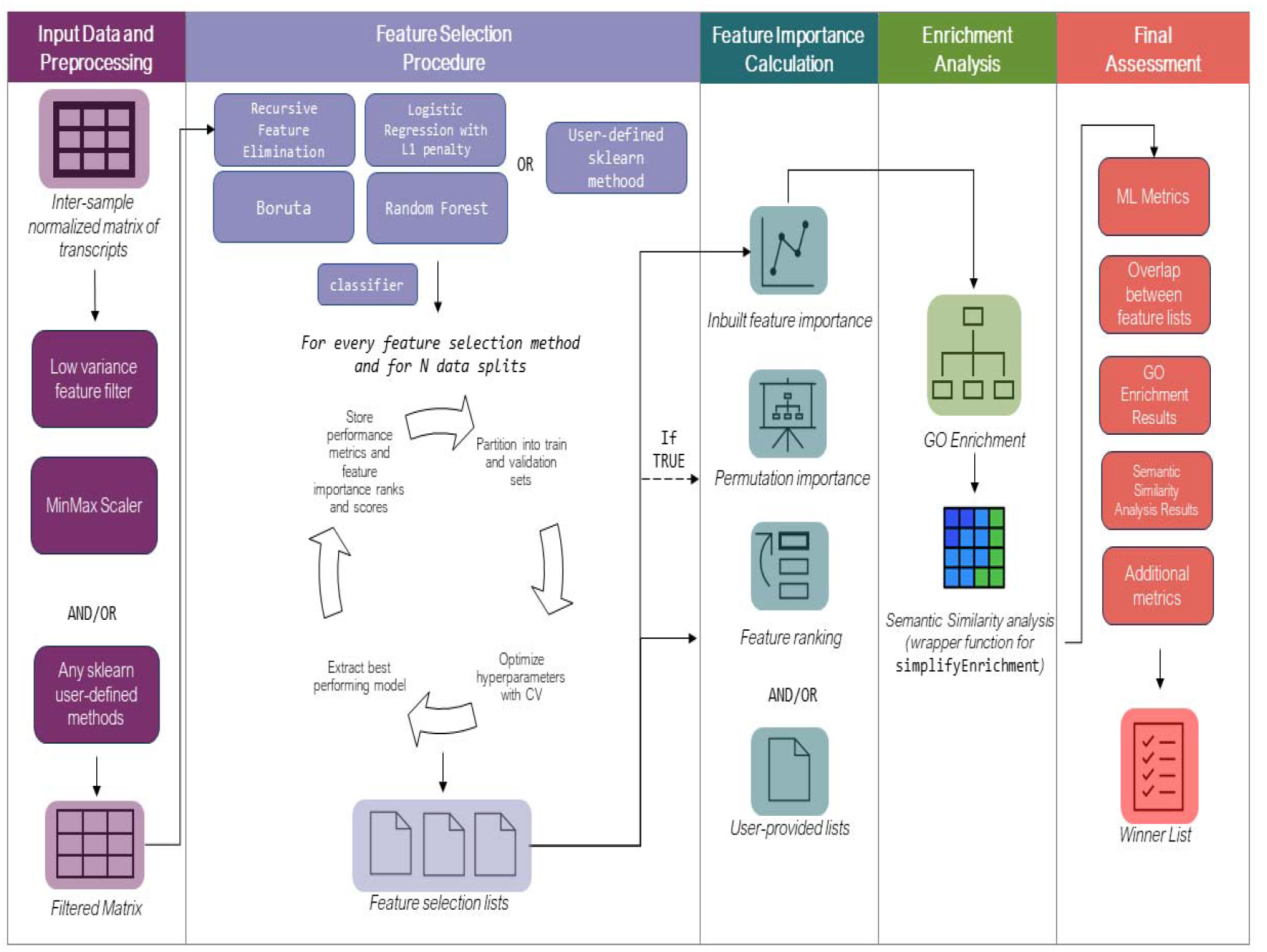
Schematic representation of the GeneSelectR package. (A) A matrix of between-sample normalized counts is subjected to initial prefiltering. (B) A feature selection procedure with four different ML methods is applied with cross validation and hyperparameter adjustment. (C) Feature importance is calculated with inbuilt feature importance and, if enabled, permutation feature importance. (D) Gene Ontology enrichment analysis is performed on every list of selected features followed by semantic similarity analysis of the significantly enriched GO terms. (E) In the final assessment, the best list is determined based on the various GeneSelectR output metrics.

Data Splitting: There is an option to keep a holdout test set, and if selected, it is possible to specify the number of random train-test splits using a customizable train-test split ratio.

Cross-validation (CV): The train set, which can either be the full dataset or a fraction of it depending on the data splitting option chosen above, is further divided into train and validation sets with a user-modifiable train-validation split ratio to perform k-fold cross-validation (CV) with hyperparameter adjustment. Users can choose from various hyperparameter search methods.

Performance Metrics: For each ML method, the best-performing pipeline is extracted, and classification performance metrics based on the CV results are calculated. If a holdout test set was chosen to be retained, performance metrics for the unseen test data are provided.

Feature Importance: Feature importance is evaluated using method-specific inbuilt feature selection. If selected, permutation feature importance is calculated.

Results Compilation: All the results are compiled into an object containing the best-performing pipelines, mean CV scores, mean test set scores, feature importance scores with ranking at every split, the selection results with inbuilt feature importance, and potentially permutation feature importance.

### 2.3 Biological Relevance Assessment with Clustering of Enriched Gene Ontology Terms

To assess the biological relevance of the different selected feature lists with regard to the biological question, a functional Gene Ontology (GO) enrichment analysis is performed, which is implemented as a wrapper function for the clusterProfiler package (Wu 2021; Thomas *et al*. 2022). In these analyses, it is also possible to include additional lists, such as results from DGE analysis. Following the GO enrichment analysis, it is possible to examine the significantly enriched GO terms within each gene list.

To arrive at a quantitative assessment on the overlap of the significantly enriched GO terms in the different features lists, a semantic similarity analysis of the GO terms is performed via a wrapper function for the simplifyEnrichment R package that takes into account the directed acyclic structure of the GO graph (Gu and Hübschmann 2023). This will lead to the clustering of enriched GO terms, displaying a heat map with the number of significant GO terms in each cluster and a word cloud for the GO terms found in each cluster. Examining these word clouds for GO terms related to the biological question will assist in selecting the feature list that contains the highest number of biologically relevant enriched GO terms assigned to the selected features.

### 2.4 Additional Assessment Functions

GeneSelectR is a highly flexible workflow that is compatible with native sklearn methods and most of the sklearn-compatible external Python libraries so that additional functions can easily be implemented. Additional functions provided within GeneSelectR with which you can examine and compare feature lists are:

1) Calculation of overlap coefficients: Based on the assumption that more robust lists of features tend to be more similar to each other, we assess their similarity by calculating overlap coefficients between the different lists using Jaccard, Soerensen-Dice and Overlap coefficient. The output here are heatmaps showing pairwise scores between every list for every method, which can provide a useful measure to assess their relevance.

2) Calculation of fractions of GO terms of interest: GO terms of interest with regard to the biological questions are expected to be associated with a number of selected transcripts in biologically relevant features lists. Given some user-defined GO terms, this function allows to quantify how many of the children nodes of the GO terms of interest can be found in the different feature lists.

### 2.5 Final Assessment

After having completed the full GeneSelectR workflow, the best-performing feature list can be determined based on 1) a high ML performance both in the CV procedure and in the test set, 2) a high overlap coefficient with the other selected feature lists, 3) a high number of biologically relevant enriched GO terms, 4) in these enriched GO terms, a high fraction of child GO terms that are part of GO parent terms of interest, and 5) a high number of GO terms associated with a relevant cluster in the semantic similarity analysis as opposed to clusters with rather broad GO terms. In addition, the size of the list should be manageable for the intended downstream analyses and data mining approaches.

## 3 Use Case

We employed GeneSelectR on a subset of 300 samples from the TCGA-BRCA dataset to identify transcriptomic biomarkers for four different breast tumor subtypes (see Supplementary Material 2 for the analysis rundown) (Parker *et al*. 2009; Weinstein *et al*. 2013). Applying the four default feature selection methods on this dataset results in marginal differences between the mean accuracy scores (mas) in the 10-fold CV (Boruta: mas: 0.921, sd: 0.012; Lasso: mas: 0.915, sd: 0.008; RandomForest: mas: 0.912, sd: 0.008; Univariate: 0.905, sd: 0.012), and in the holdout test data (Boruta: mas: 0.876, sd: 0.027; RandomForest: mas: 0.876, sd: 0.036; Univariate: 0.863, sd: 0.015; Lasso: mas: 0.858, sd: 0.039). The range of the number of selected features across the four ML algorithms varied from 10 for Univariate to 106, as identified by Boruta. In contrast, the list of differentially expressed genes was much longer with 1826 genes that both had an adjusted p-value smaller than 0.001 and a fold change higher than 5. The Overlap coefficient (OC) for the overlap between the feature lists based on permutation feature importance is highest for DEGs compared to the Univariate list (OC: 0.65). The second-highest OC of 0.5 is achieved when comparing the Lasso-selected list to both the Boruta-selected and the RandomForest-selected lists. These criteria based on ML performance and comparisons between the different lists do not provide enough information to make a well-founded selection of the best list.

As an additional criterion for the selection of the best-performing list, the biological relevance of the different selected feature lists is assessed in GeneSelectR using an overrepresentation analysis to identify enriched GO Biological Process (BP) terms. This revealed that angiogenesis, glycosaminoglycan metabolism, and the apoptotic p53 pathway, which are implicated in crucial cancer-related processes, were significantly enriched in the Boruta-selected feature list. For a comparison of the GO BP terms that are enriched in the different lists taking into account the GO graph, a semantic similarity analysis using Louvain clustering was performed, which grouped the GO BP terms into six distinct clusters. Notably, two of these clusters focused on glucosamine metabolic process and humoral immune response. These were enriched in the Boruta-selected list, further substantiating its biological relevance. In addition, the number of child nodes for the two GO terms *cell cycle regulation* and *immune response* were quantified among the enriched GO terms in the different lists. The Boruta-selected feature list showed a high fraction of child nodes for both of these GO terms, again demonstrating its relevance in cancer-related processes.

Overall, Boruta was identified as the best-performing feature selection method. In comparison, DGE analysis yielded a broader, less interpretable list of differentially expressed genes, emphasizing its limitations in analyzing complex datasets.

## 4 Conclusion

We introduce here GeneSelectR, a novel workflow that synergistically integrates the ML capabilities of the sklearn Python package with the bioinformatics functionalities of R libraries like clusterProfiler and simplifyEnrichment. This approach addresses the limitations of traditional DGE analysis in selecting differential features in complex datasets by enabling the capture of nonlinear relationships between genes. GeneSelectR offers a robust feature selection procedure that not only optimizes ML performance, but also ensures biological relevance through functional enrichment and semantic similarity analyses.

Our application of GeneSelectR to the TCGA-BRCA dataset demonstrated its efficacy in identifying a list of transcriptomic biomarkers that are relevant to the biomedical question. The Boruta feature selection method, in particular, provided a list that was superior in ML classification performance and more biologically relevant compared to the other ML methods. Additionally, it was more specific and maintained a manageable size compared to the DGE analysis list. The workflow’s flexibility and compatibility with external libraries make it a versatile tool for various bioinformatics applications.

GeneSelectR thus presents a significant advancement in the field, offering a comprehensive solution for gene selection that is both computationally efficient and biologically insightful.

## Supporting information

Supplementary Material: TCGA-BRCA Example

## Acknowledgements

We would like to thank the Swiss Institute of Allergy and Asthma Research and the Center for Data Analysis, Visualization and Simulation (DAViS) at the University of Applied Sciences of the Grisons for providing resources to conduct the project.

