## Supplementary Table 1 for "GeneSelectR: An R Package Workflow for Enhanced Feature Selection from RNA Sequencing Data"

| Method/param | threshold | mode | param | C | solver | penalty | perc | n_estimators | max_depth | min_samples_split | min_samples_leaf | bootstrap | max_features |
|---|---|---|---|---|---|---|---|---|---|---|---|---|---|
| StandardScaler | X | X | X | X | X | X | X | X | X | X | X | X | X |
| VarianceThreshold | 0.85 | X | X | X | X | X | X | X | X | X | X | X | X |
| Feature Selector:Univariate | X | k_best | [50,100, 150,200] | X | X | X | X | X | X | X | X | X | X |
| Feature Selector: Lasso | X | X | X | [0.01, 0.1, 1, 10] | [liblinear, saga] | l1 | X | X | X | X | X | X | X |
| Feature Selector: boruta | X | X | X | X | X | X | [80,90,100] | [50,100,250, 500] | X | X | X | X | X |
| Feature Selector: RandomForest | X | X | X | X | X | X | X | [100,150,200,250,300,350,400,450,500] | [3,10,20,30] | [2,5,10] | [1,2,4] | [True,False] | X |
| Classifier: RandomForest | X | X | X | X | X | X | X | [100,500,1000] | [3,10,20,30,40,50] | [2,5,10] | [1,2,4] | X | [auto,sqrt] |

Supplementary Table 1. Default Parameters of Feature Selection Methods
