## Supplementary figures and images for "GeneSelectR: An R Package Workflow for Enhanced Feature Selection from RNA Sequencing Data"

### Supplementary Figure 1

#### GeneSelectR Algorithm Flowchart

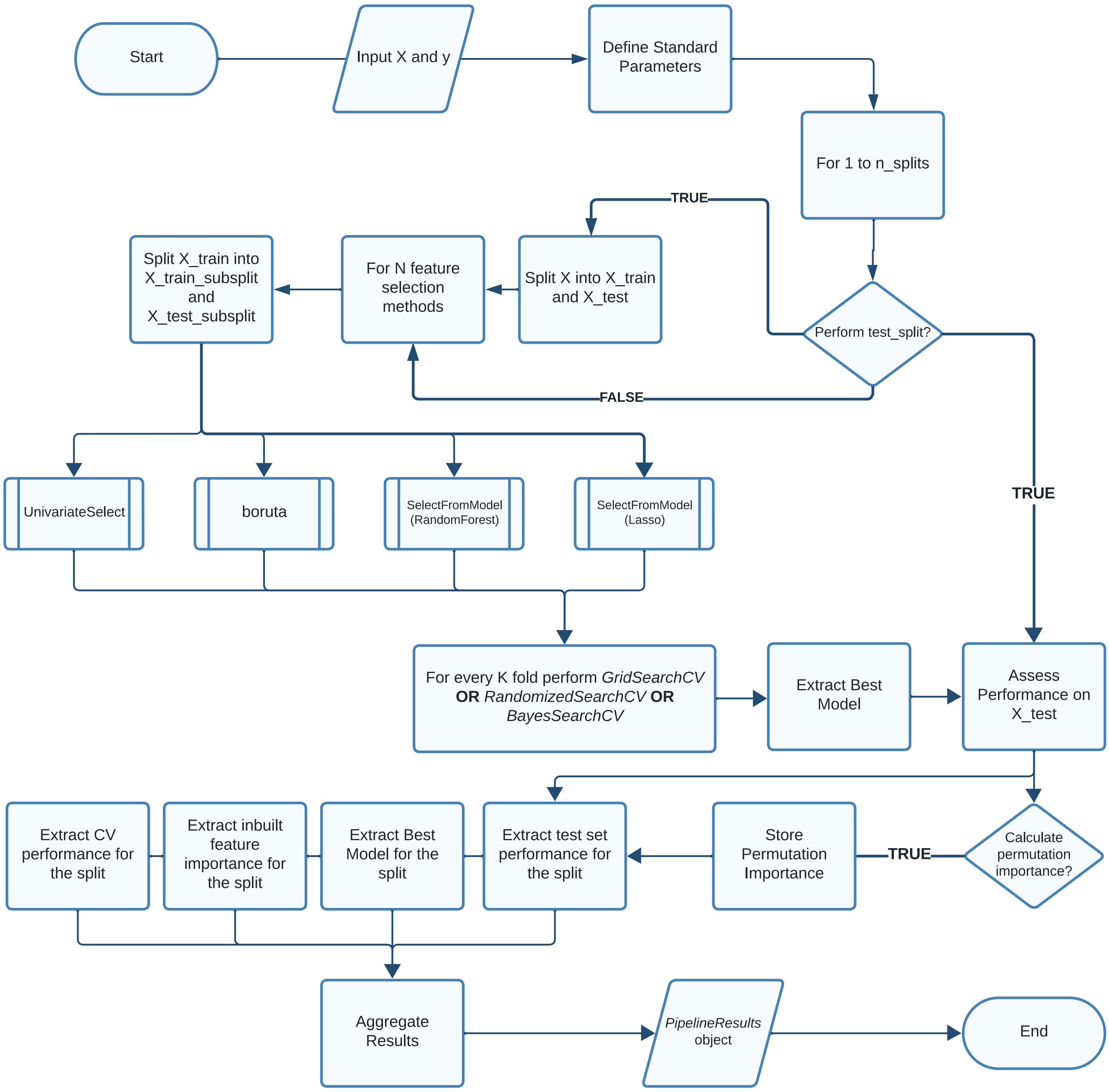
