## Supplementary Material: TCGA-BRCA Example for "GeneSelectR: An R Package Workflow for Enhanced Feature Selection from RNA Sequencing Data"

### GeneSelectR Application on TCGA-BRCA Dataset

Damir Zhakparov

2023-10-19

#### Introduction

This tutorial walks you through the use of `GeneSelectR` with the **TCGA-BRCA RNA Expression dataset** from The Cancer Genome Atlas. To acquire the dataset, we employed the `TCGAbiolinks` R package. For a step-by-step guide on data extraction, consult this script.

##### Biological Question

The primary aim of this tutorial is to identify a transcriptomic signature that is specific for the four subtypes of primary breast cancer tumor defined here by their PAM50 markers.

By doing so, we hope to shed light on the unique molecular profile of each subtype, which could subsequently inform targeted therapeutic strategies and prognostic assessments.

#### Dataset Overview

In this guide, we focus on a subset of **380 samples**, all categorized under the **'Primary Solid Tumor'** type and labeled with the four tumor subtypes : Basal(n = 80), Her2(n = 38), LumA(n = 188), and LumB (n = 74). The analysis encompasses all **60,600 sequenced transcripts**. For an in-depth look at the TCGA-BRCA dataset, please visit the official documentation. The data files used in the tutorial can be accessed in this repository.

#### Molecular Subtypes Overview

##### Basal-like (basal)

- **Characteristics**: Strong expression of basal markers (cytokeratines), high levels of Ki-67
- **Prognosis**: Poor
- **Standard Treatment**: Chemotherapy

##### HER2-enriched (Her2)

- **Characteristics**: HER2 Positive (HER2+), Hormone Receptor Negative (HR-)
- **Prognosis**: Intermediate
- **Standard Treatment**: HER2-targeted therapies with the monoclonal antibody trastuzumab

##### Luminal A (LumA)

- **Characteristics**: Hormone Receptor Positive (HR+), low levels of HER2 and Ki-67
- **Prognosis**: Best among the subtypes
- **Standard Treatment**: Hormone therapy

##### Luminal B (LumB)

- **Characteristics**: Hormone Receptor Positive (HR+), higher levels of HER2 and Ki-67
- **Prognosis**: Worse than Luminal A, but better than Basal-like
- **Standard Treatment**: Hormone therapy, may require chemotherapy

By understanding the unique transcriptomic landscape of each subtype, we can better predict disease outcomes and tailor treatment regimens.

### 1. Differential Gene Expression Analysis

As a baseline differential gene expression (DGE) analysis provides a good starting point. First of all, we will load the dataset and metadata files and import the necessary packages:

```r
# set up working directories
data_dir <- file.path('../raw-data')
output_dir <- file.path('../results')
```

```r
#load the files
sample_metadata <- readRDS(file.path(data_dir, 'sample_metadata.rds'))
raw_counts <- readRDS(file.path(data_dir, 'raw_counts.rds'))
```

After that we will create a DESEq2 object and then filter out genes with low counts:

```r
dds <- DESeqDataSetFromMatrix(countData = raw_counts,
                              colData = sample_metadata,
                              design = ~ paper_BRCA_Subtype_PAM50)

# filter the low counts
table(sample_metadata$paper_BRCA_Subtype_PAM50)
#>
#> Basal  Her2  LumA  LumB
#>    80    38   188    74
smallestGroupSize <- 38 #smallest group is 38 samples
keep <- rowSums(counts(dds) >= 10) >= smallestGroupSize
dds <- dds[keep,]
```

Before performing the DGE analysis it's strongly advised to perform PCA on normalized counts to see if there any batch effects:

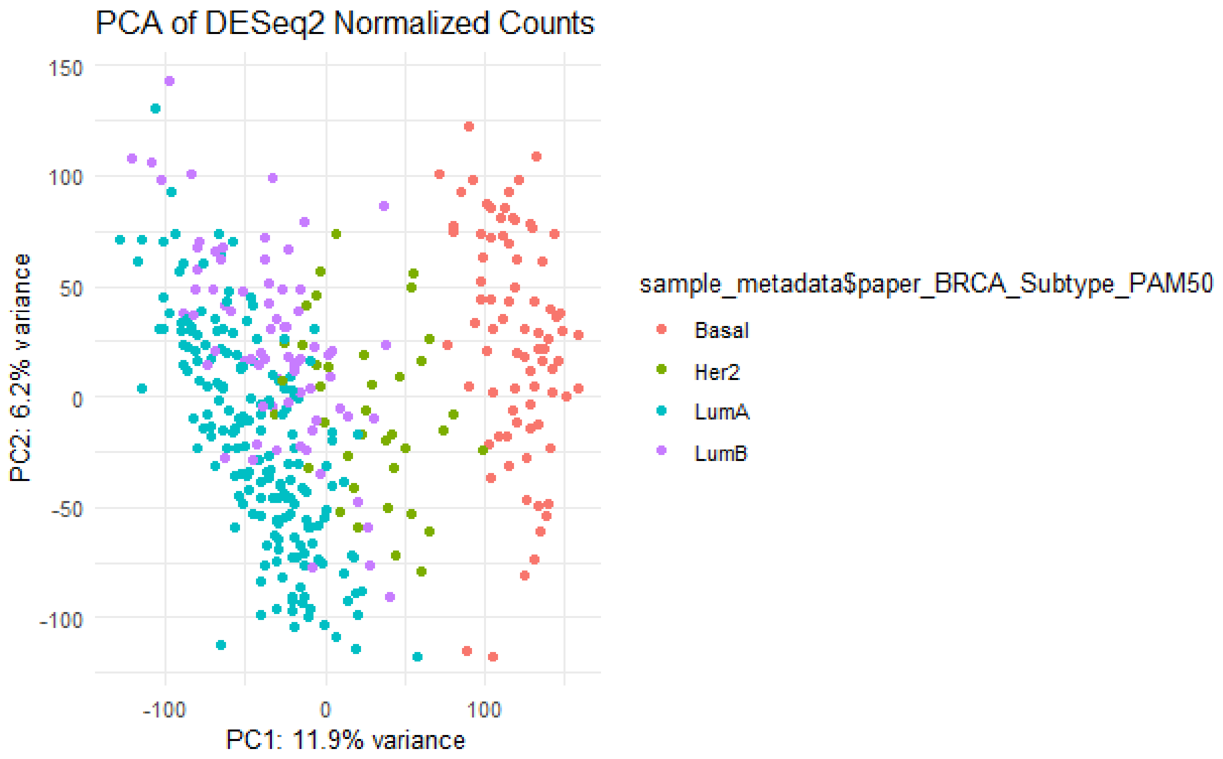

If there are factors in the metadata that account for major sources of variation in your data, these should be included in the design formula. And now let's run the DGE analysis:

```
# Run DESeq
dds <- DESeq(dds)
```

After the calculations are done, let's filter out differentially expressed genes:

```
# Get the DGE results
res <- results(dds)
filtered_res <- subset(res, padj < 0.001 & abs(log2FoldChange) > log2(5))
filtered_df <- as.data.frame(filtered_res)
```

And store it in a separate vector:

```
deg_list <- rownames(filtered_df)
# remove the splice information from the gene names
deg_list <- gsub("\\.\\d+", "", deg_list)
```

#### 2. Feature Selection with GeneSelectR

##### 2.1 Data Preparation

GeneSelectR expects the data to be a matrix/data.frame where rows represent samples and columns are features. Please note that the dataset has to be normalized within samples prior to any analysis with the package. So now we extract the vst-transformed matrix from our DESeq2 object for feature selection:

```
# Extract VST (Variance-Stabilized Transformed) Data
vsd <- vst(dds, blind = FALSE)
vsd_matrix <- assay(vsd)
vsd_matrix <- t(vsd_matrix)
```

Then we will create a response vector y with sample labels:

```
sample_metadata <-
sample_metadata[match(rownames(vsd_matrix),rownames(sample_metadata)), ]
sample_metadata$paper_BRCA_Subtype_PAM50 <-
as.factor(sample_metadata$paper_BRCA_Subtype_PAM50)
sample_metadata$num_label <-
as.integer(sample_metadata$paper_BRCA_Subtype_PAM50) # NOTE: the labels
should be encoded with numeric values
table(sample_metadata$num_label)
#>
#>   1   2   3   4
#>  80  38 188  74
```

##### 2.2 Running GeneSelectR

After preparing the data we are ready to run GeneSelectR:

```
X <-  exp
y <- sample_metadata %>% select(num_label)
```

```
selection_results <- GeneSelectR(X, # gene expression matrix
                                 y, # label vector
                                 njobs = -1, # number of cores to be deployed
(-1 = all)
                                 n_splits = 5, # number of train/test splits
to perform
                                 max_features = 250, # max features to be
selected per feature selection method (only RF, Lasso and Univariate)
                                 perform_test_split = TRUE, # if partition of
the data into train and test
                                 scoring = 'accuracy', # scoring metric for
optimization
                                 calculate_permutation_importance = TRUE, #
whether to calculate permutation importance
                                 search_type = 'random', # type of grid
search
                                 n_iter = 150L) # number of hyperparameter
combinations to be sampled
```

The ENSEMBL IDs contain the splice variant number  at the end following a dot, which is not convenient for further analyses. We can remove it by running:

```
convert_ensembl_to_symbol <- function(selection_results) {
  target_slots <- c("inbuilt_feature_importance", "permutation_importance")

  for (slot_name in target_slots) {
    slot_content <- slot(selection_results, slot_name)

    if (!is.null(slot_content)) {
      new_slot_content <- lapply(names(slot_content), function(method) {
        df <- slot_content[[method]]

        # Remove digits after the period in feature column
        df$feature <- gsub("\\.\\d+", "", df$feature)

        # Convert Ensembl IDs to gene symbols
        converted <- clusterProfiler::bitr(df$feature, fromType = "ENSEMBL",
toType = "SYMBOL", OrgDb = 'org.Hs.eg.db')

        # Left join to preserve all original rows
        return(merge(df, converted, by.x = "feature", by.y = "ENSEMBL", all.x
= TRUE))
      })

      names(new_slot_content) <- names(slot_content)
      slot(selection_results, slot_name) <- new_slot_content
    }
  }
  return(selection_results)
```

```
}

# Apply the function
selection_results <- convert_ensembl_to_symbol(selection_results)
```

#### 3. Analyzing the Results

After the analysis has been finished, we can look into the PipelineResults objects to inspect the results. For example if we call:

```
str(selection_results, max.level = 2)
#> Formal class 'PipelineResults' [package "GeneSelectR"] with 6 slots
#>   ..@ best_pipeline              :List of 5
#>   ..@ cv_results                 :List of 5
#>   ..@ inbuilt_feature_importance:List of 4
#>   ..@ permutation_importance    :List of 4
#>   ..@ cv_mean_score              :'data.frame':   4 obs. of  3 variables:
#>   ..@ test_metrics               : tibble [4 × 9] (S3: 
tbl_df/tbl/data.frame)
```

We can see the following slots in the Pipeline results object:

- best_pipeline - contains all the parameters for the best performing pipeline

- cv_results - contains the entire output of the CV procedure during hyperparameter search

- inbuilt_feature_importance - inbuilt feature importance with mean, std and rank for every feature across iterations
- permutation_importance - permutation feature importance with mean, std and rank for every feature across iterations;
- cv_mean_scores - CV scores but aggregated into one dataframe with mean and std of the optimization metrics scores;
- test_metrics - metrics (f1, recall, precision and accuracy) for the unseen test set. Let's inspect some of the most interesting metrics for the evaluation.

##### 3.1 Machine Learning Performance Metrics

First, let's explore the cross validation scores for each of our methods. We can inspect the CV mean performance by displaying the data.frame, which by default gives the accuracy scores:

```
selection_results@cv_mean_score
#>          method mean_score    sd_score
#> 1        boruta  0.9210374 0.012047342
#> 2         Lasso  0.9153061 0.007984372
#> 3 RandomForest  0.9127721 0.007926976
#> 4    Univariate  0.9045578 0.011999833
```

We can see that Boruta is slightly better in terms of CV performance, but all methods are somewhat comparable. In this case, we can inspect the performance on unseen data stored in the test_metrics slot:

```
selection_results@test_metrics
#> # A tibble: 4 × 9
#>   method       f1_mean  f1_sd recall_mean recall_sd precision_mean
precision_sd
#>   <chr>          <dbl>  <dbl>       <dbl>     <dbl>          <dbl>
<dbl>
#> 1 Lasso          0.854 0.0351       0.858    0.0388          0.870
0.0262
#> 2 RandomForest   0.873 0.0346       0.876    0.0356          0.883
0.0286
#> 3 Univariate     0.859 0.0127       0.863    0.0150          0.866
0.0136
#> 4 boruta         0.876 0.0251       0.876    0.0273          0.889
0.0234
#> # ℹ 2 more variables: accuracy_mean <dbl>, accuracy_sd <dbl>
```

Again, Boruta seems to be the best-performing method, although marginally. We can produce a combined plot of all metrics by calling:

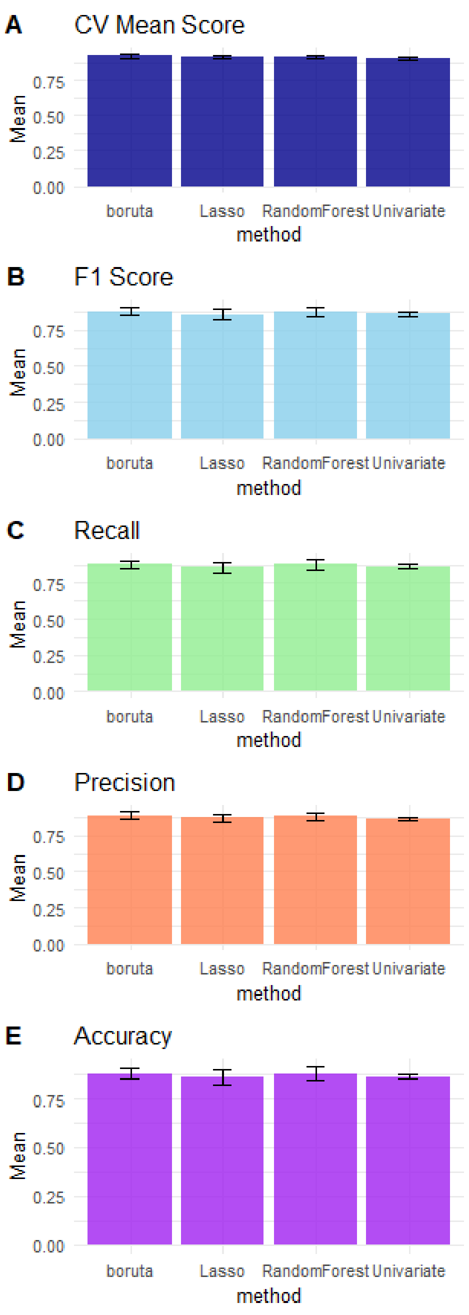

#### 3.2 Feature Importance

The next step is to inspect the most important features for each feature selection method. To plot the importance scores, we will call the '**plot_feature_importance()' function** This function returns a list of plots demonstrating mean feature importance scores across different data splits, which we will store in a separate object:

```
plot_list <- plot_feature_importance(selection_results)
plot_list
#> $Lasso
```

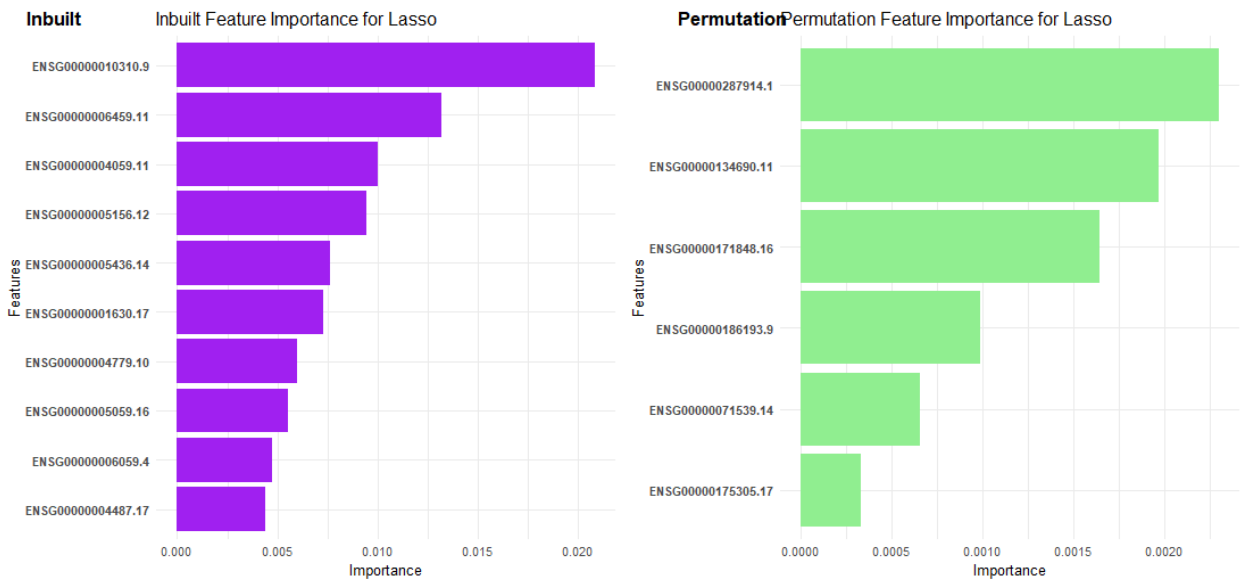

```
#>
#> $Univariate
```

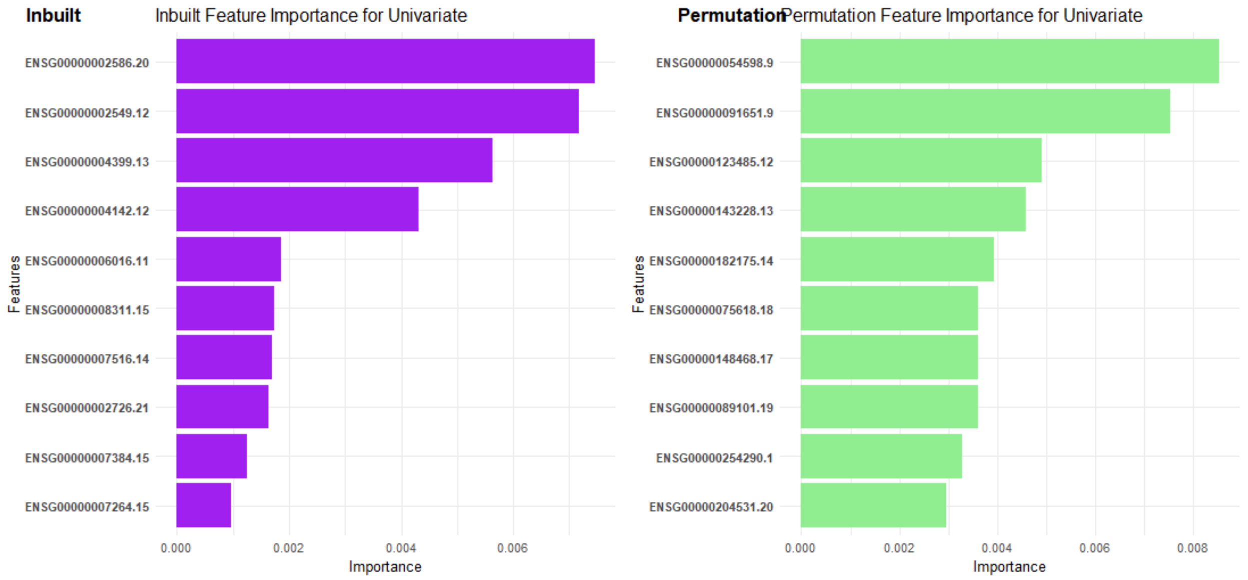

```
#>
#> $RandomForest
```

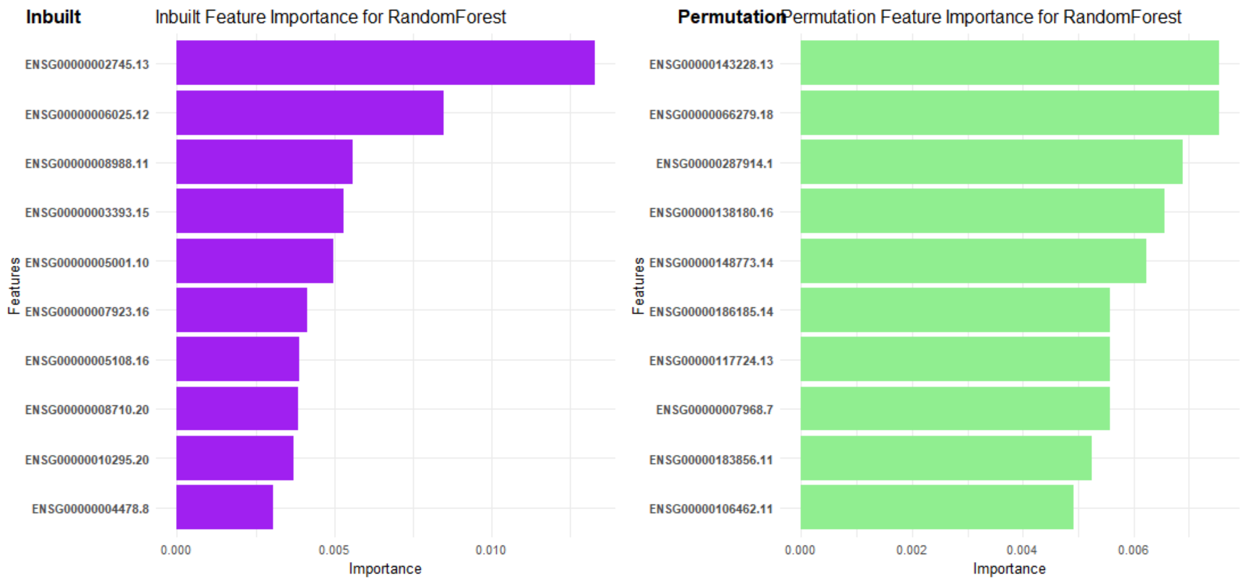

```
#>
#> $boruta
```

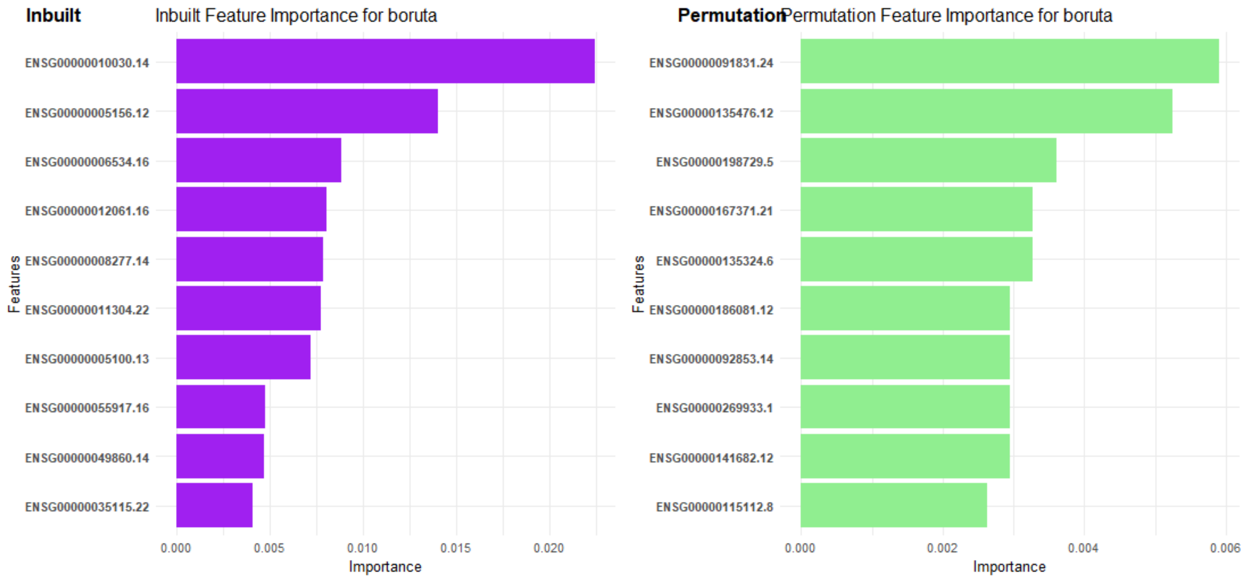

Interestingly, we can see that Boruta has a lot of relevant genes to cancer as top features, e.g.: ENSG00000010030 (ETV7), ENSG00000005156 (LIG3), ENSG00000006534 (ALDH3B1), ENSG00000005100 (DHX33).

#### 3.3 Overlap between Gene Lists and DGE list

It might be interesting to see if there is any overlap between the feature selection lists also including the list of differentially expressed genes (DEGs).

Then we can calculate the overlap coefficients and plot them:

```
#inspect overlap with DEGs
overlap_degs <- calculate_overlap_coefficients(selection_results,
```

```
custom_lists = list('DEGS' = deg_list))
plot_overlap_heatmaps(overlap_degs)
```

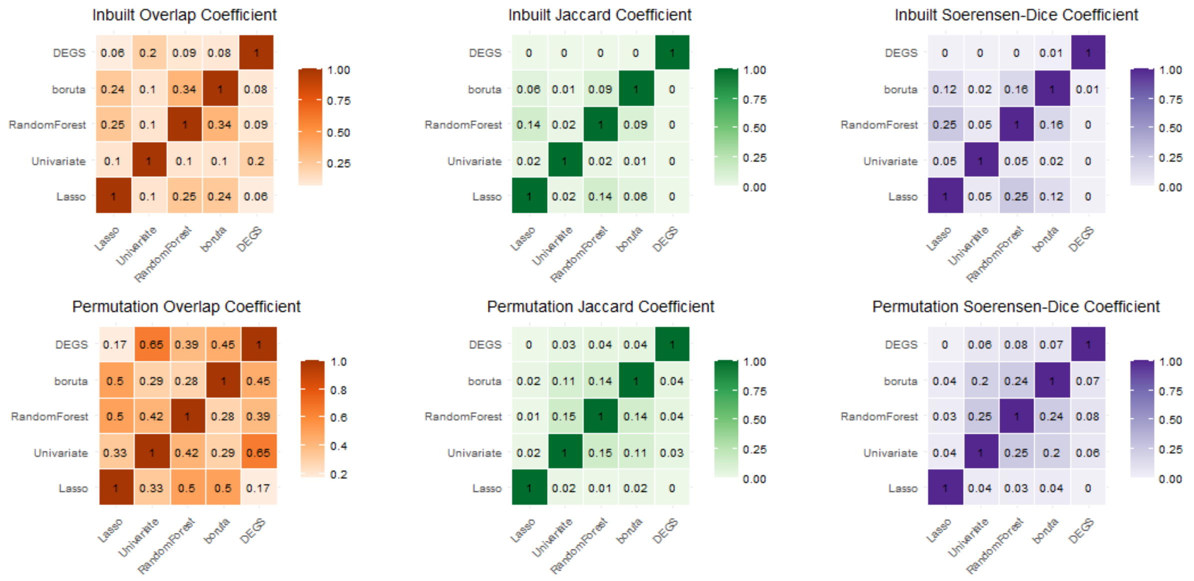

We can see three different coefficients that calculate the list a bit differently. Since our lists are different in sizes, the most relevant one here is Overlap Coefficient. We can see that some of the permutation importance lists have similarities. For example, Lasso is quite similar to Boruta and RandomForest. As for the DEGs list it's quite similar to the Univariate method, which is somewhat expected.

We can also inspect whether there is an overlap with the canonical PAM50 signature. To do so we can use the following code:

```
# load the PAM50 signature
pam50 <- read.csv(file = file.path(data_dir, 'PAM50.txt'), sep = '\t', header
= FALSE, col.names = c('SYMBOL'))

# convert to ensembl ids
pam50_ens <- clusterProfiler::bitr(pam50$SYMBOL, fromType = "SYMBOL", toType
= "ENSEMBL", OrgDb = 'org.Hs.eg.db')
#> 'select()' returned 1:many mapping between keys and columns

#calculate the overlap with the PAM50 signature
overlap_pam50 <- calculate_overlap_coefficients(selection_results,
custom_lists = list('PAM50' = pam50_ens$ENSEMBL, 'DEGS' = deg_list))
plot_overlap_heatmaps(overlap_pam50)
```

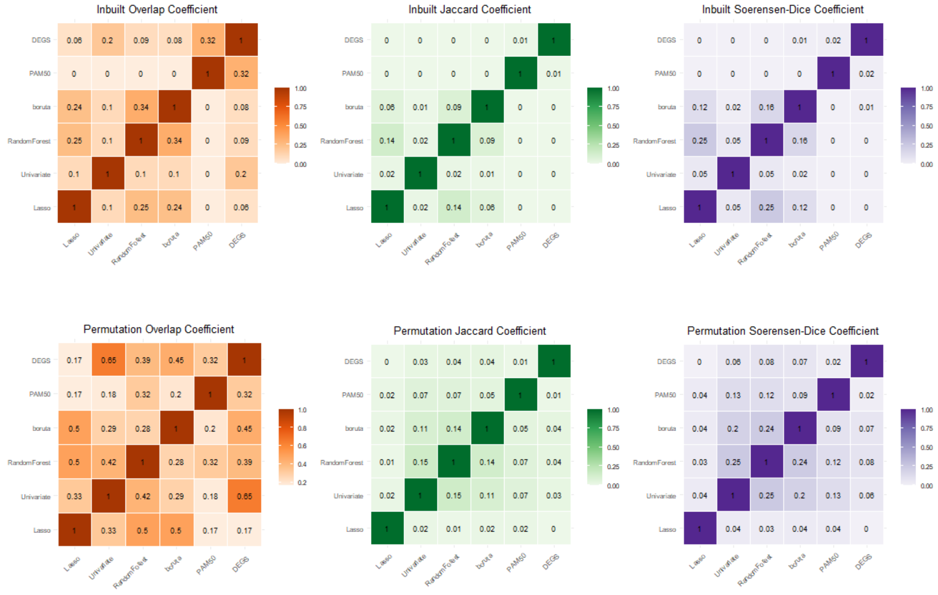

Finally, we can see the exact numbers of overlapping features by producing an UpSet plot:

```r
plot_upset(selection_results, custom_lists = list('PAM50' =
pam50_ens$ENSEMBL, 'DEGS' = deg_list))
#> $inbuilt_importance
```

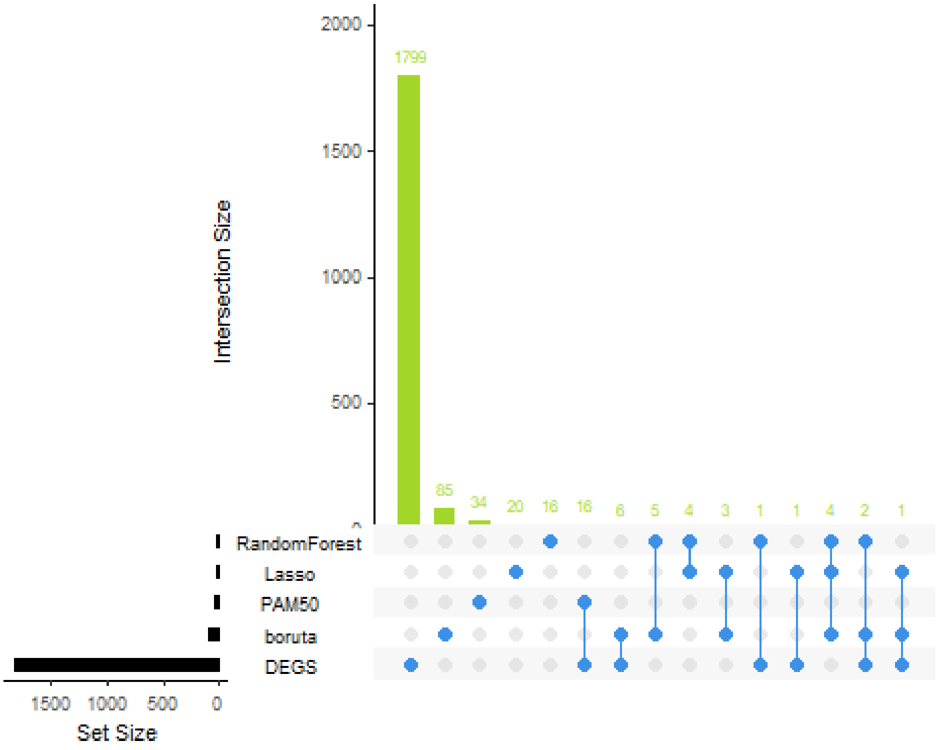

```r
#>
#> $permutation_importance
```

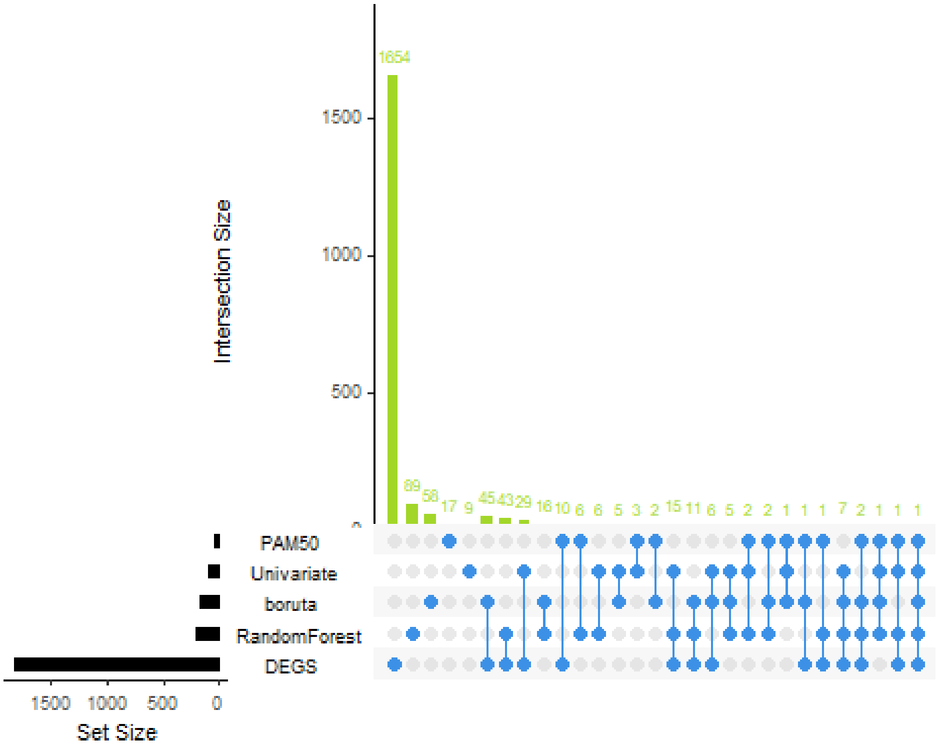

There is some overlap of the PAM50 signature with the DEGs, but no overlap with any of the feature selection lists based on inbuilt feature importance. As for the permutation importance, there is some overlap with the PAM50 signature.

#### 3.4 Gene List Annotations

For the enrichment analyses and subsequent biological interpretation it is convenient to convert gene identifiers to others that are appropriate/useful in different situations. To do so we can do the following:

```
# remove the splice variant number from the ensembl ids
background <- as.character(colnames(vsd_matrix))
background <- gsub("\\..*", "", background)
custom_list <- list('background' = background,
                    'DEGs' = deg_list)

ah <- AnnotationHub::AnnotationHub()

# connect to the annotation database “EnsDb” and then fetch an annotation for the human genome, which is ‘AH98047’
human_ens <- AnnotationHub::query(ah, c("Homo sapiens", "EnsDb"))
human_ens <- human_ens[['AH98047']]
#> loading from cache
#> require("ensembldb")
annotations_ahb <- ensembldb::genes(human_ens, return.type = "data.frame")
%>%
```

```
  dplyr::select(gene_id,gene_name,entrezid,gene_biotype)

annotations_df <- annotate_gene_lists(pipeline_results = selection_results,
                                      annotations_ahb = annotations_ahb,
                                      format = 'ENSEMBL',
                                      custom_lists = custom_list)
#> 'select()' returned 1:1 mapping between keys and columns
#> 'select()' returned 1:1 mapping between keys and columns
#> 'select()' returned 1:1 mapping between keys and columns
#> 'select()' returned 1:1 mapping between keys and columns
#> 'select()' returned 1:many mapping between keys and columns
#> 'select()' returned 1:many mapping between keys and columns
#> 'select()' returned 1:1 mapping between keys and columns
#> 'select()' returned 1:1 mapping between keys and columns
#> 'select()' returned 1:1 mapping between keys and columns
#> 'select()' returned 1:many mapping between keys and columns
#> 'select()' returned 1:many mapping between keys and columns
#> 'select()' returned 1:many mapping between keys and columns
```

This returns an object of class AnnotatedGeneLists containing gene symbol, ENSEMBL ID and ENTREZ ID for the selected features as well as background and DEG lists.

#### 3.5 Gene Ontology Enrichment

Now we will do Gene Ontology Enrichment analysis to mine the pathways relevant to our biological question. To do so we can call the function:

```
# perform GO Analysis
annotated_GO_inbuilt <- GO_enrichment_analysis(annotations_df,
                                       list_type = 'inbuilt', #run GO
enrichment on inbuilt  selected features
                                       keyType = 'ENSEMBL', # run analysis
with ENSEMBLIDs
                                       background = background,
                                       ont = 'BP') # run BP ontology
#> Performing GO Enrichment analysis for the:Lasso
#> Performing GO Enrichment analysis for the:Univariate
#> Performing GO Enrichment analysis for the:RandomForest
#> Performing GO Enrichment analysis for the:boruta
#> Performing GO Enrichment analysis for the:DEGs

annotated_GO_permutation <- GO_enrichment_analysis(annotations_df,
                                           list_type = 'permutation',
#run GO enrichment on permutation based selected features
                                           keyType = 'ENSEMBL', # run
analysis with ENSEMBLIDs
                                           background = background,
                                           ont = 'BP') # run BP ontology
#> Performing GO Enrichment analysis for the:Lasso
#> Performing GO Enrichment analysis for the:Univariate
```

```
#> Performing GO Enrichment analysis for the:RandomForest
#> Performing GO Enrichment analysis for the:boruta
#> Performing GO Enrichment analysis for the:DEGs
```

So far, inbuilt feature importance looked promising, so now we can inspect what GO terms are enriched in each gene list:

```
# code snippet to plot the GO terms

ggplot(data = annotated_GO_inbuilt$RandomForest@result %>%
         arrange(pvalue) %>%
         head(10) %>%
         mutate(GeneCount = as.numeric(gsub("/.*$", "", GeneRatio))),
       aes(x = reorder(Description, -log10(pvalue)), y = -log10(pvalue))) +
  geom_point(aes(size = GeneCount, color = pvalue)) +
  scale_color_gradient(low = "red", high = "blue") +
  scale_size_continuous(range = c(3, 9)) +
  #geom_text(aes(label = GeneRatio), vjust = -1) +
  coord_flip() +
  xlab("Top 10 GO Term Descriptions") +
  ylab("-log10(pvalue)") +
  ggtitle('Univariate List') +
  theme_minimal()
```

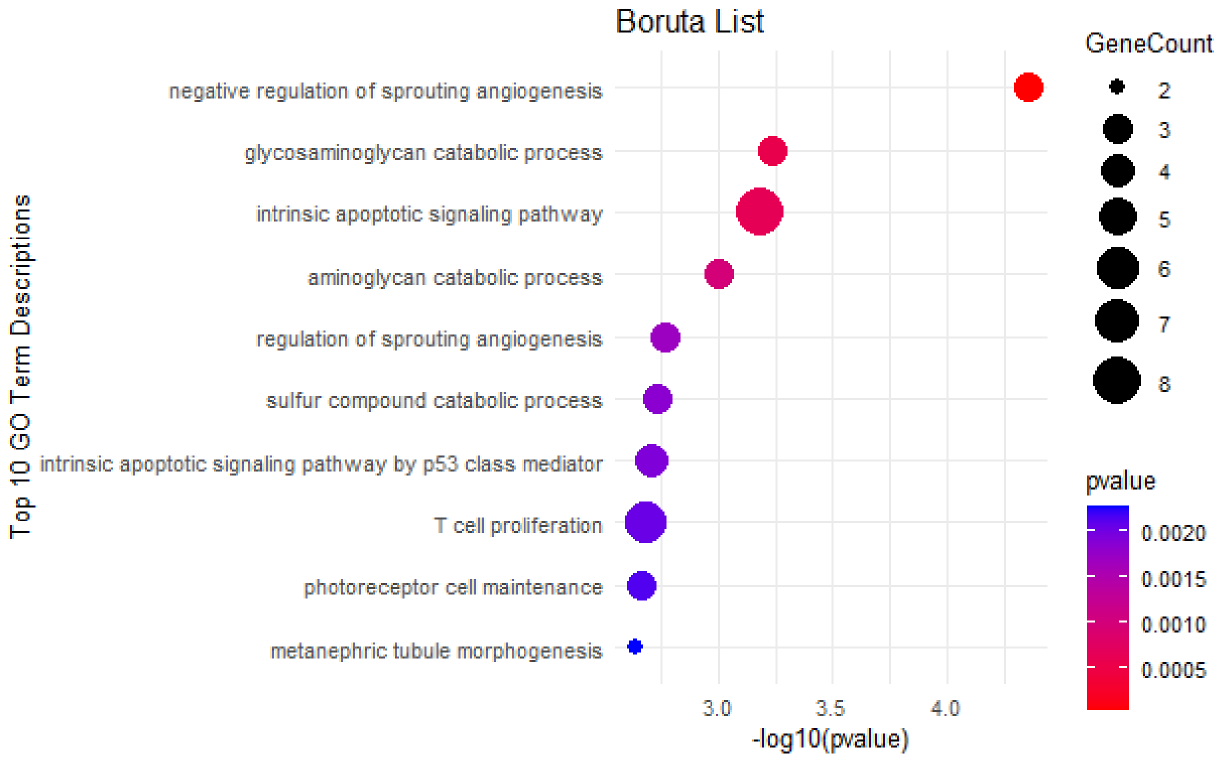

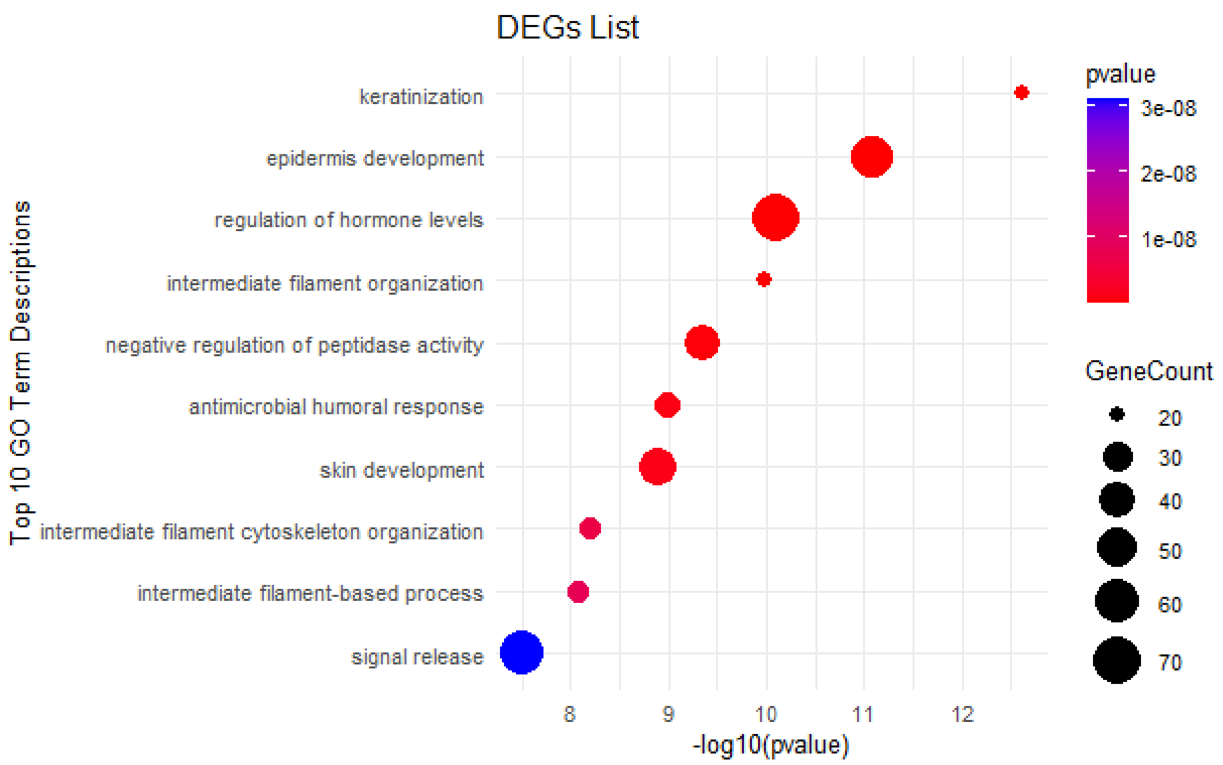
DEGs List
Top 10 GO Term Descriptions
keratinization
epidermis development
regulation of hormone levels
intermediate filament organization
negative regulation of peptidase activity
antimicrobial humoral response
skin development
intermediate filament cytoskeleton organization
intermediate filament-based process
signal release
8
9
10
11
12
-log10(pvalue)
pvalue
3e-08
2e-08
1e-08
GeneCount
20
30
40
50
60
70

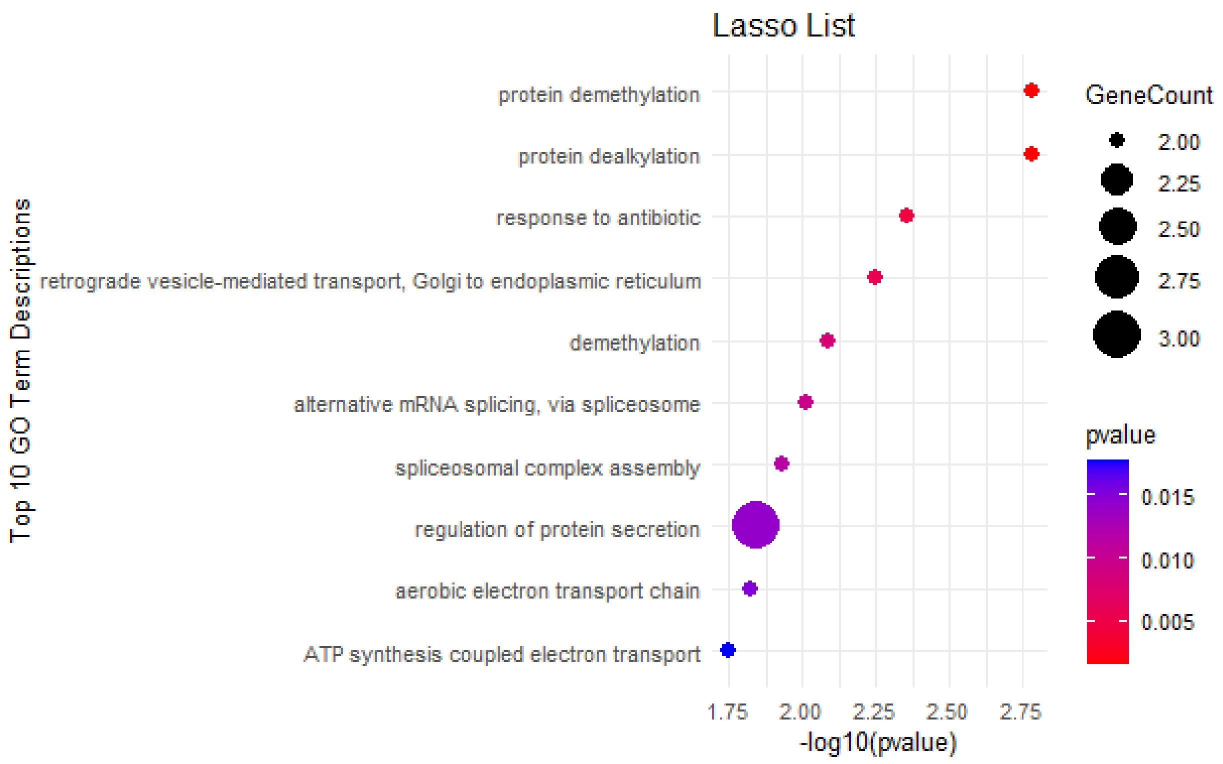
Lasso List
Top 10 GO Term Descriptions
protein demethylation
protein dealkylation
response to antibiotic
retrograde vesicle-mediated transport, Golgi to endoplasmic reticulum
demethylation
alternative mRNA splicing, via spliceosome
spliceosomal complex assembly
regulation of protein secretion
aerobic electron transport chain
ATP synthesis coupled electron transport
1.75
2.00
2.25
2.50
2.75
-log10(pvalue)
GeneCount
2.00
2.25
2.50
2.75
3.00
pvalue
0.015
0.010
0.005

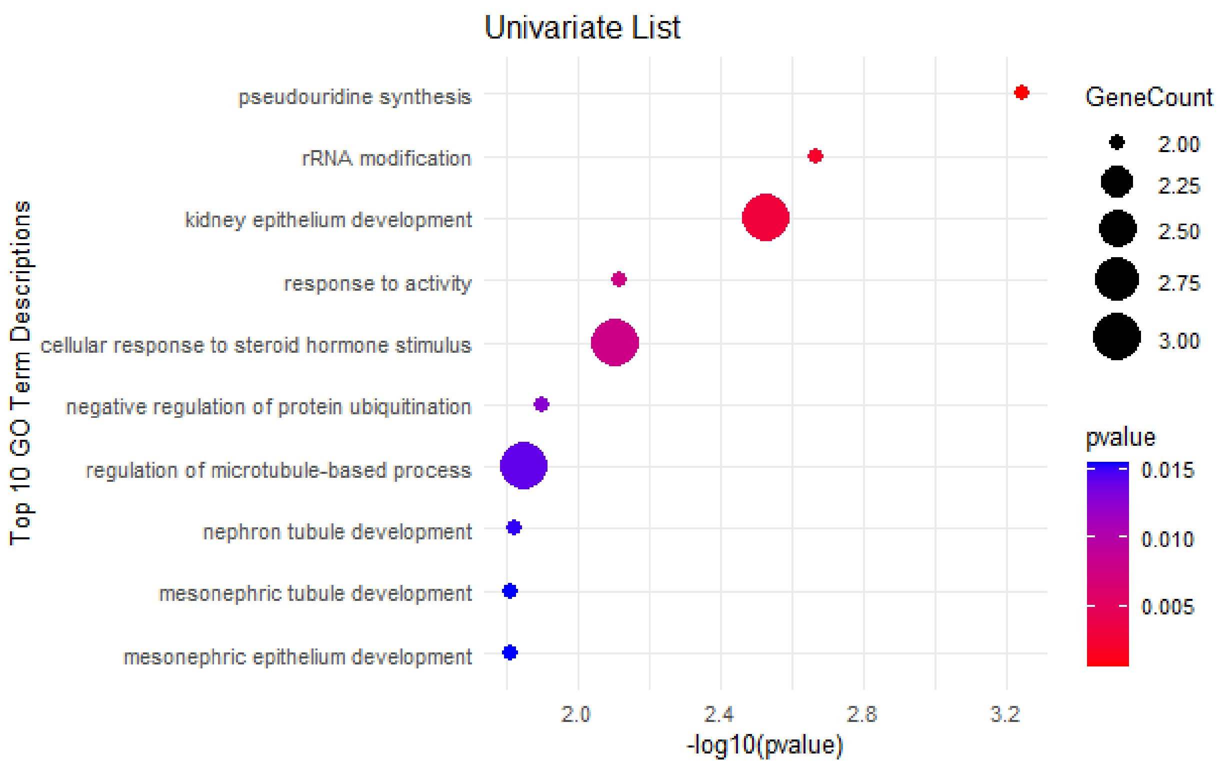

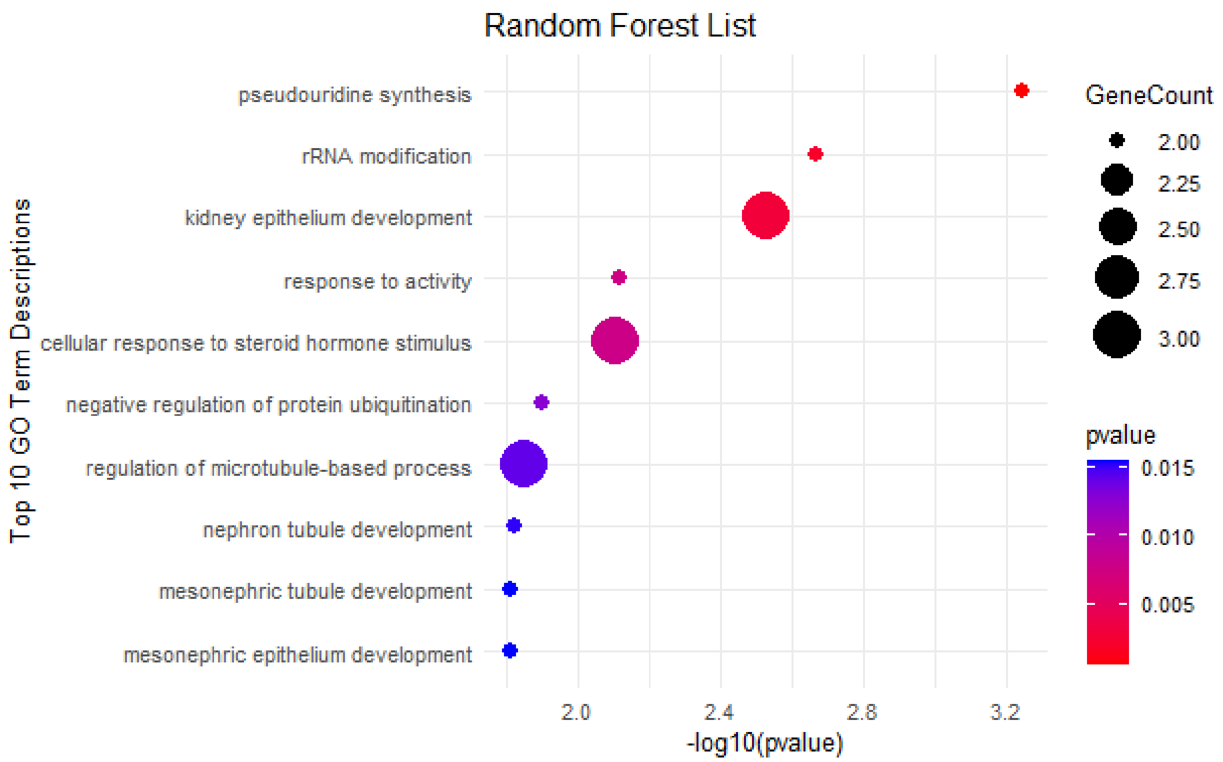

When examining the top 10 enriched pathways, it becomes apparent that Boruta identifies the most relevant ones. For example, we can see terms related to the regulation of sprouting angiogenesis, which are associated with metastasis spread. Additionally, terms such as glycosaminoglycan metabolic process and especially apoptotic p53 pathway are all of high relevance in relation to cancer.

##### 3.6 Quantification of Children Nodes of a Parent Node of Interest

Additionally, we can quantify how many of the children nodes of a parent node of interest are in our list. For example, we can take two relevant broad GO Biological Process (BP) terms that are *cell cycle regulation* (GO:0051726) and *immune response* (GO:0006955).

```
annot_child_fractions_inbuilt <- compute_GO_child_term_metrics(GO_data =
annotated_GO_inbuilt,
                              GO_terms = c("GO:0051726", "GO:0006955"),
                              plot = TRUE)
```

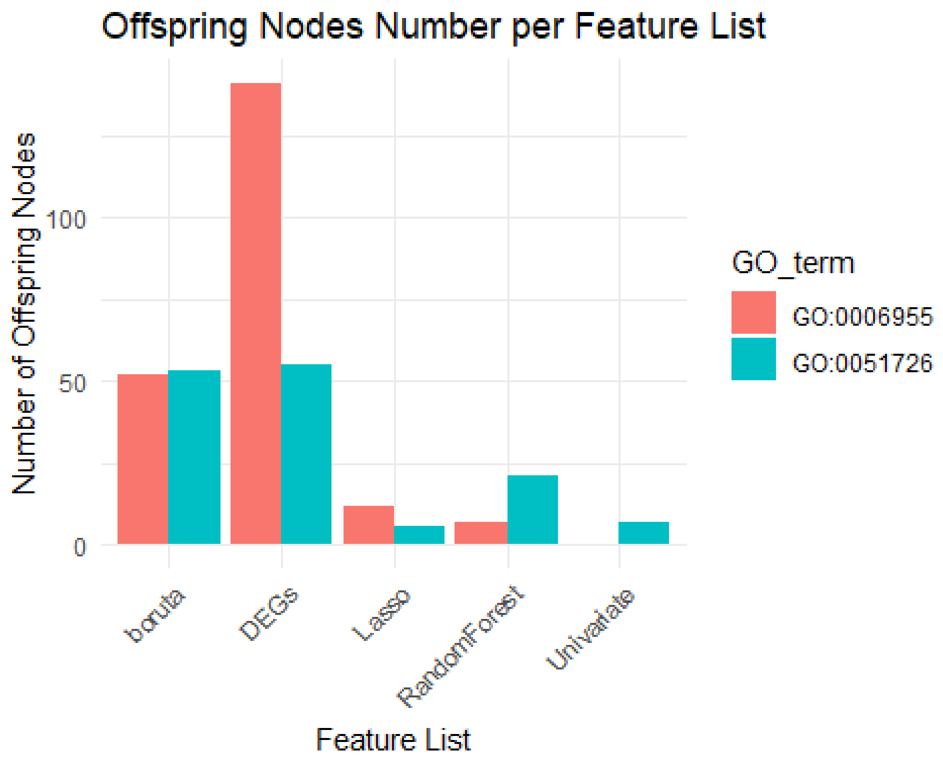

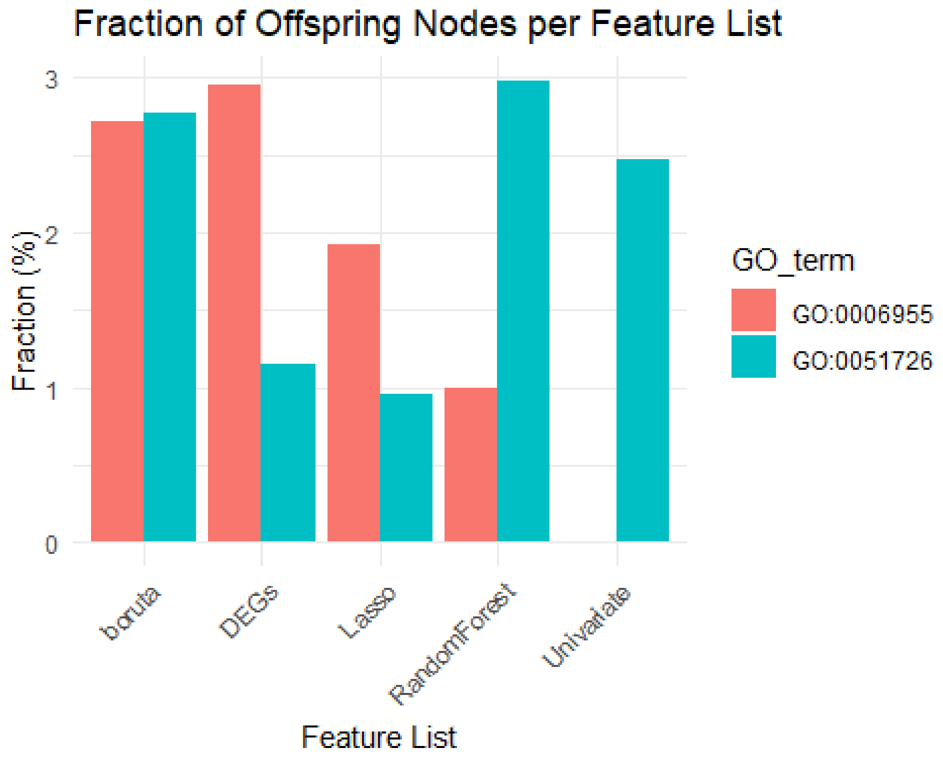

```
annot_child_fractions_permut <- compute_GO_child_term_metrics(GO_data =
annotated_GO_permutation,
```

```
                              GO_terms = c("GO:0051726", "GO:0006955"),
                              plot = TRUE)
```

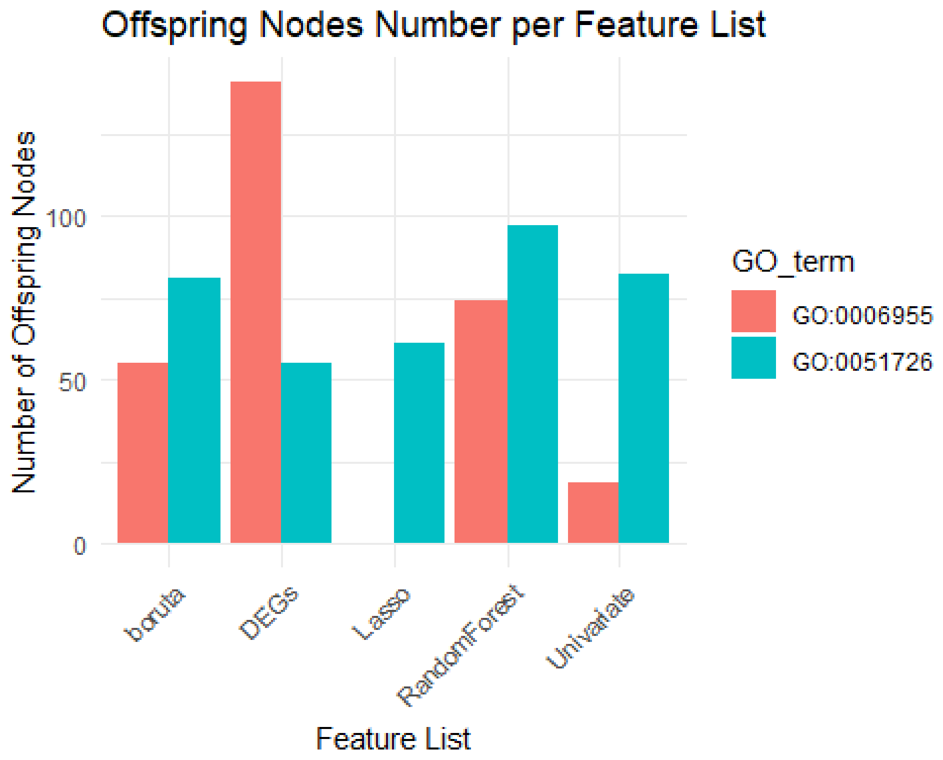

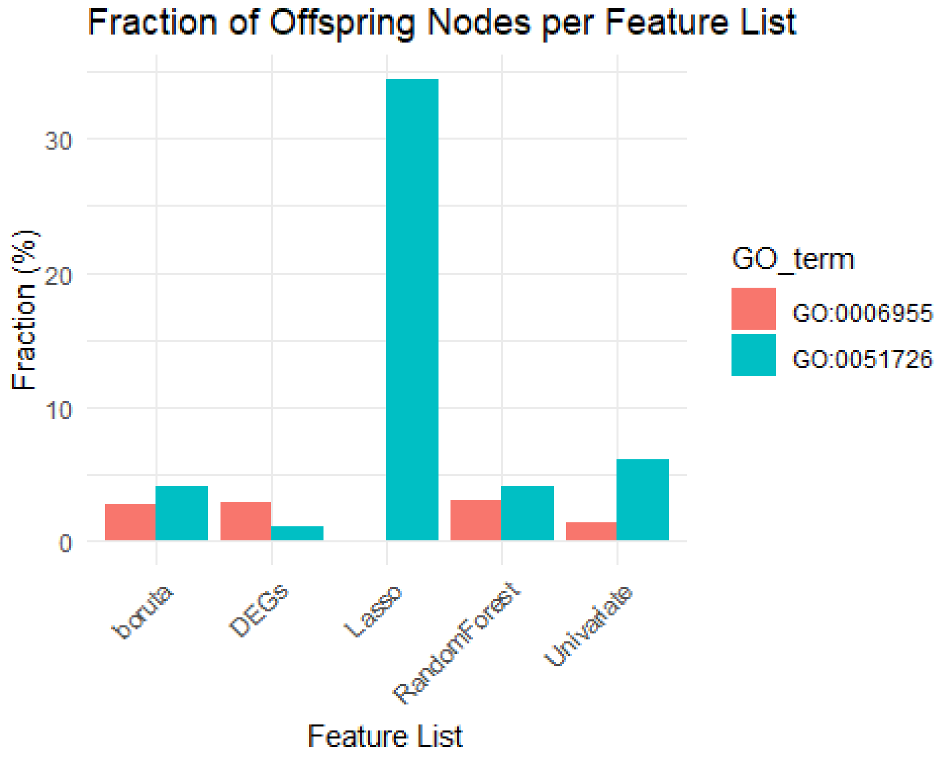

Once again, we see that Boruta performs very well in terms of fractions of the parent terms of interest *immune process* and *cell cycle regulation*.

##### 3.7 Semantic Similarity Analysis

Finally, we can now perform semantic similarity analysis of identified GO terms. To do that we will run this:

```
# perform the semantic similarity analysis
hmap_inbuilt <- run_simplify_enrichment(annotated_GO_inbuilt,
                                method = 'louvain',
                                measure = 'Rel',
                                ont = 'BP',
                                padj_column = 'pvalue',
                                padj_cutoff = 0.01)
#> Use column 'ID' as `go_id_column`.
#> Loading required namespace: gridtext
#> 470/5074 GO IDs left for clustering.
#> Cluster 470 terms by 'louvain'... 7 clusters, used 0.09124589 secs.
#> 'magick' package is suggested to install to give better rasterization.
#>
#> Set `ht_opt$message = FALSE` to turn off this message.
#> Perform keywords enrichment for 6 GO lists...
#> 'magick' package is suggested to install to give better rasterization.
#>
#> Set `ht_opt$message = FALSE` to turn off this message.
```

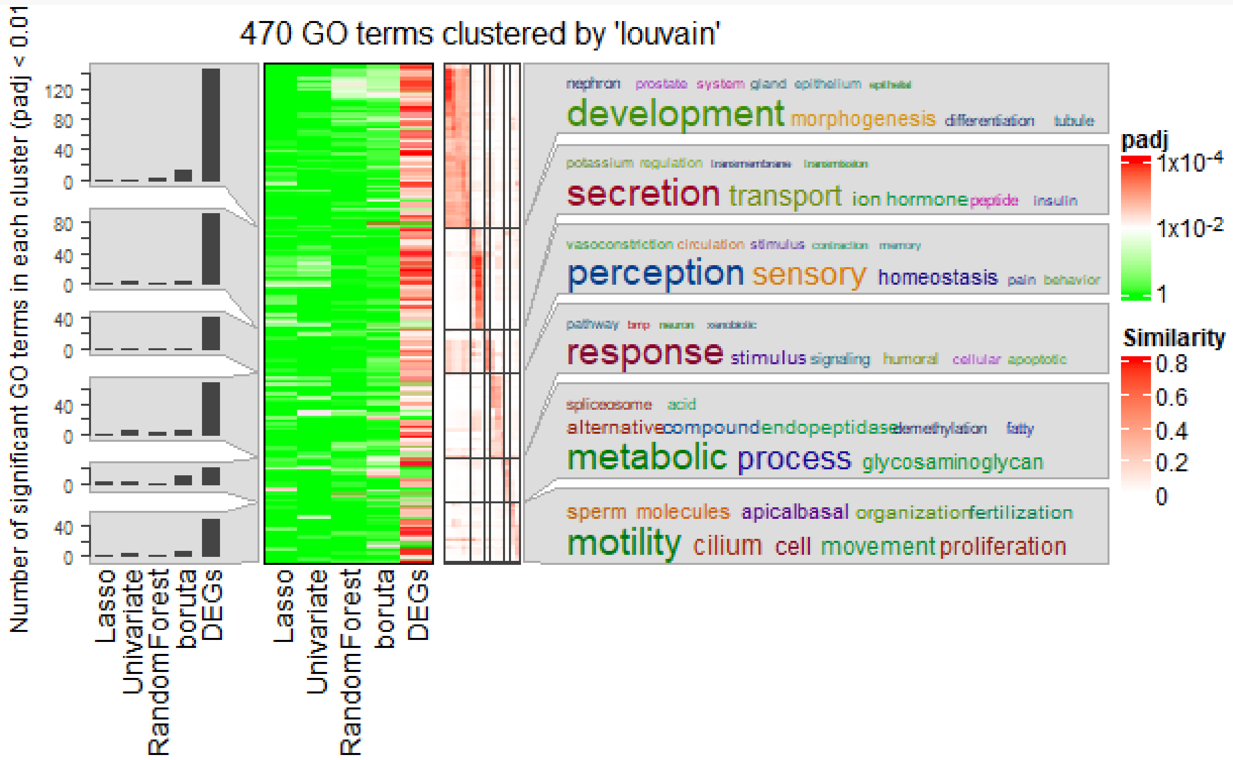

```
hmap_permutation <- run_simplify_enrichment(annotated_GO_permutation,
                                            method = 'louvain',
                                            measure = 'Jiang',
                                            ont = 'BP',
                                            padj_column = 'pvalue',
                                            padj_cutoff = 0.01)
#> Use column 'ID' as `go_id_column`.
#> 470/5074 GO IDs left for clustering.
#> Cluster 470 terms by 'louvain'... 9 clusters, used 0.04489207 secs.
#> 'magick' package is suggested to install to give better rasterization.
#>
#> Set `ht_opt$message = FALSE` to turn off this message.
#> Perform keywords enrichment for 7 GO lists...
#> 'magick' package is suggested to install to give better rasterization.
#>
#> Set `ht_opt$message = FALSE` to turn off this message.
```

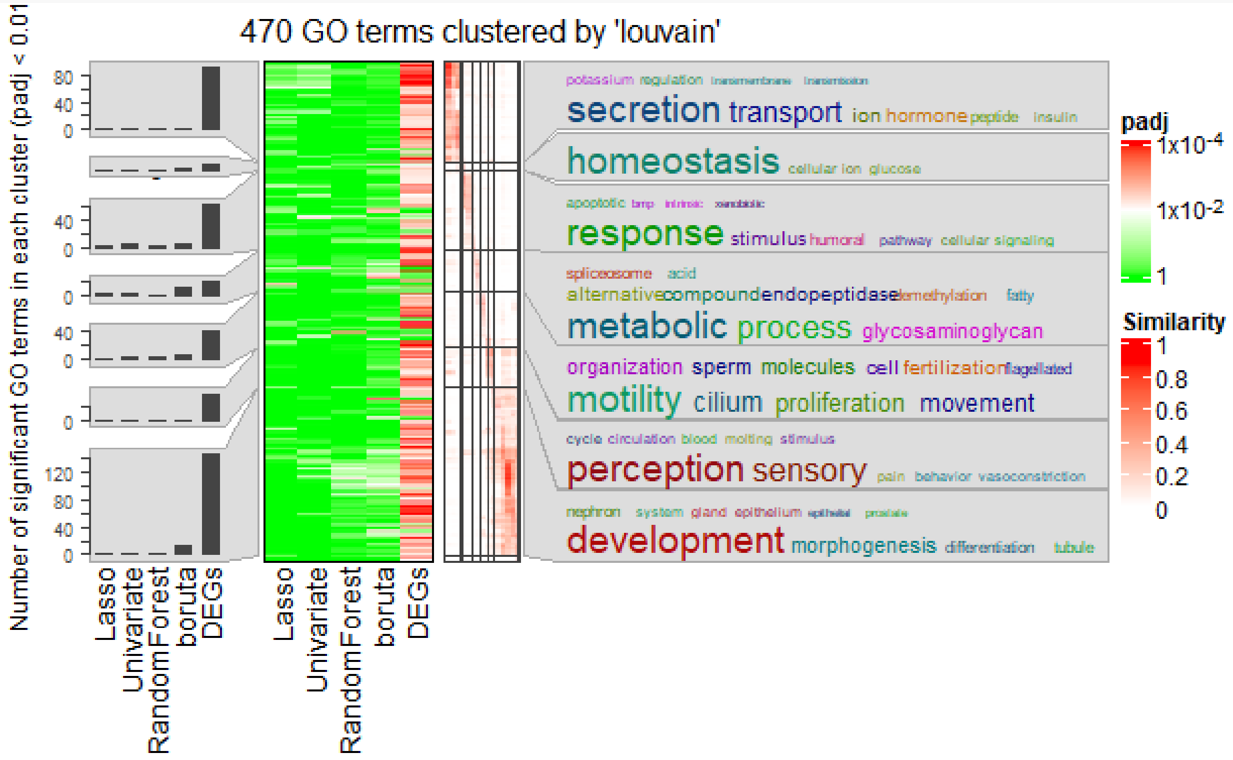

After performing the semantic similarity analysis, we observe 6 clusters in total. By examining word clouds, we can identify two clusters of interest: cluster 4 contains GO terms related to humoral immunity and cell signal transduction, and cluster 5 contains GO terms related to glycosaminoglycan metabolism and other metabolism-related terms. If we examine the significance and fraction heatmap on the left, we can observe that the GO terms enriched in the list of DEGs is significant across the board, without displaying any specificity. In contrast, GO terms enriched in the Boruta list are significantly enriched in these two clusters.

#### 4. Picking the Winner and Conclusions

After completing the analysis, we can now choose the winner among the lists. From different points of view Boruta seems to be the best-performing list because: 1. It has the highest ML performance in the CV and test sets; 2. The Boruta list gives more significantly enriched GO BP terms related to differentiation between molecular subtypes that are related to cancer cell metabolism, apoptosis and angiogenesis; 3. the Boruta list has higher fractions of child terms that are part of the parent terms of interest in comparison to other lists;
4. In terms of semantic similarity analysis, the Boruta list is more specific to GO terms related to immune response and metabolism, as compared to the list of DEGs that results in rather broad and unspecific GO BP terms; and 5. In comparison to the list of DEGs, the Boruta list is much shorter and more manageable for downstream analyses and interpretation of the results. With this, the feature list selected by Boruta seems to be the best candidate for downstream analyses and targeted validation experiments to establish some of the features as biomarkers. Based on this, we conclude that Boruta performed best in selecting biomedically relevant features in the TCGA-BRCA dataset..

```
utils::sessionInfo()
#> R version 4.3.0 (2023-04-21 ucrt)
#> Platform: x86_64-w64-mingw32/x64 (64-bit)
#> Running under: Windows 11 x64 (build 22621)
#> 
#> Matrix products: default
#> 
#> 
#> locale:
#> [1] LC_COLLATE=English_Switzerland.utf8  LC_CTYPE=English_Switzerland.utf8
#> [3] LC_MONETARY=English_Switzerland.utf8 LC_NUMERIC=C
#> [5] LC_TIME=English_Switzerland.utf8
#> 
#> time zone: Europe/Zurich
#> tzcode source: internal
#> 
#> attached base packages:
#> [1] stats4    stats     graphics  grDevices utils     datasets  methods
#> [8] base
#> 
#> other attached packages:
#>  [1] ensembldb_2.24.0            AnnotationFilter_1.24.0
#>  [3] GenomicFeatures_1.52.2      AnnotationDbi_1.62.2
#>  [5] ggplot2_3.4.4               DESeq2_1.40.2
#>  [7] SummarizedExperiment_1.30.2 Biobase_2.60.0
#>  [9] MatrixGenerics_1.12.3       matrixStats_1.0.0
#> [11] GenomicRanges_1.52.0        GenomeInfoDb_1.36.2
#> [13] IRanges_2.34.1              S4Vectors_0.38.1
#> [15] BiocGenerics_0.46.0         GeneSelectR_0.0.0.9000
#> [17] dplyr_1.1.2
#> 
```

```
#> loaded via a namespace (and not attached):
#>   [1] ProtGenerics_1.32.0           bitops_1.0-7
#>   [3] enrichplot_1.20.1             HDO.db_0.99.1
#>   [5] httr_1.4.7                    RColorBrewer_1.1-3
#>   [7] doParallel_1.0.17             tools_4.3.0
#>   [9] utf8_1.2.3                    R6_2.5.1
#>  [11] lazyeval_0.2.2                GetoptLong_1.0.5
#>  [13] withr_2.5.1                   prettyunits_1.2.0
#>  [15] gridExtra_2.3                 plotwidgets_0.5.1
#>  [17] cli_3.6.1                     scatterpie_0.2.1
#>  [19] labeling_0.4.3                slam_0.1-50
#>  [21] tm_0.7-11                     commonmark_1.9.0
#>  [23] Rsamtools_2.16.0              yulab.utils_0.0.9
#>  [25] gson_0.1.0                    DOSE_3.26.1
#>  [27] rstudioapi_0.15.0             RSQLite_2.3.1
#>  [29] generics_0.1.3                gridGraphics_0.5-1
#>  [31] shape_1.4.6                   BiocIO_1.10.0
#>  [33] gtools_3.9.4                  GO.db_3.17.0
#>  [35] Matrix_1.6-1                  fansi_1.0.4
#>  [37] abind_1.4-5                   lifecycle_1.0.3
#>  [39] yaml_2.3.7                    gplots_3.1.3
#>  [41] qvalue_2.32.0                 BiocFileCache_2.8.0
#>  [43] grid_4.3.0                    blob_1.2.4
#>  [45] promises_1.2.1                crayon_1.5.2
#>  [47] lattice_0.21-8                cowplot_1.1.1
#>  [49] KEGGREST_1.40.0               pillar_1.9.0
#>  [51] knitr_1.43                    ComplexHeatmap_2.16.0
#>  [53] fgsea_1.26.0                  rjson_0.2.21
#>  [55] codetools_0.2-19              fastmatch_1.1-4
#>  [57] glue_1.6.2                    downloader_0.4
#>  [59] ggfun_0.1.2                   data.table_1.14.8
#>  [61] vctrs_0.6.3                   png_0.1-8
#>  [63] treeio_1.24.3                 testthat_3.2.0
#>  [65] gtable_0.3.4                  cachem_1.0.8
#>  [67] xfun_0.40                     S4Arrays_1.0.6
#>  [69] mime_0.12                     tidygraph_1.2.3
#>  [71] pheatmap_1.0.12               iterators_1.0.14
#>  [73] interactiveDisplayBase_1.38.0 ellipsis_0.3.2
#>  [75] nlme_3.1-162                  simplifyEnrichment_1.10.0
#>  [77] ggtree_3.8.2                  bit64_4.0.5
#>  [79] progress_1.2.2                filelock_1.0.2
#>  [81] UpSetR_1.4.0                  KernSmooth_2.23-20
#>  [83] colorspace_2.1-0              DBI_1.1.3
#>  [85] tidyselect_1.2.0              proxyC_0.3.3
#>  [87] bit_4.0.5                     compiler_4.3.0
#>  [89] curl_5.0.2                    xml2_1.3.5
#>  [91] NLP_0.2-1                     DelayedArray_0.26.7
#>  [93] shadowtext_0.1.2              rtracklayer_1.60.1
#>  [95] scales_1.2.1                  caTools_1.18.2
#>  [97] rappdirs_0.3.3                stringr_1.5.0
```

```
#>  [99] digest_0.6.33                 tmod_0.50.13
#> [101] rmarkdown_2.24                XVector_0.40.0
#> [103] htmltools_0.5.6               pkgconfig_2.0.3
#> [105] highr_0.10                    dbplyr_2.3.3
#> [107] fastmap_1.1.1                 rlang_1.1.1
#> [109] GlobalOptions_0.1.2           shiny_1.7.5
#> [111] farver_2.1.1                  jsonlite_1.8.7
#> [113] BiocParallel_1.34.2           GOSemSim_2.26.1
#> [115] RCurl_1.98-1.12               magrittr_2.0.3
#> [117] GenomeInfoDbData_1.2.10       ggplotify_0.1.2
#> [119] patchwork_1.1.3               munsell_0.5.0
#> [121] Rcpp_1.0.10                   ape_5.7-1
#> [123] viridis_0.6.4                 reticulate_1.34.0
#> [125] stringi_1.7.12                tagcloud_0.6
#> [127] ggraph_2.1.0                  brio_1.1.3
#> [129] zlibbioc_1.46.0               MASS_7.3-58.4
#> [131] AnnotationHub_3.8.0           plyr_1.8.8
#> [133] org.Hs.eg.db_3.17.0           parallel_4.3.0
#> [135] ggrepel_0.9.3                 Biostrings_2.68.1
#> [137] graphlayouts_1.0.0            splines_4.3.0
#> [139] gridtext_0.1.5                hms_1.1.3
#> [141] circlize_0.4.15               locfit_1.5-9.8
#> [143] igraph_1.4.3                  markdown_1.8
#> [145] reshape2_1.4.4                biomaRt_2.56.1
#> [147] BiocVersion_3.17.1            XML_3.99-0.14
#> [149] evaluate_0.22                 RcppParallel_5.1.7
#> [151] BiocManager_1.30.22           foreach_1.5.2
#> [153] tweenr_2.0.2                  httpuv_1.6.11
#> [155] tidyr_1.3.0                   purrr_1.0.2
#> [157] polyclip_1.10-4               clue_0.3-64
#> [159] ggforce_0.4.1                 xtable_1.8-4
#> [161] restfulr_0.0.15               tidytree_0.4.5
#> [163] later_1.3.1                   viridisLite_0.4.2
#> [165] tibble_3.2.1                  clusterProfiler_4.8.3
#> [167] aplot_0.2.0                   memoise_2.0.1
#> [169] beeswarm_0.4.0                GenomicAlignments_1.36.0
#> [171] cluster_2.1.4
```
